# Viral DNA SIP: the isotopic composition of virions is correlated with that of the substrate used for host growth

**DOI:** 10.1101/2022.10.15.512377

**Authors:** Vuong Quoc Hoang Ngo, Maximilien Sotomski, Angeline Guenne, Mahendra Mariadassou, Mart Krupovic, François Enault, Ariane Bize

## Abstract

DNA Stable Isotope Probing is emerging as a powerful tool to study host-virus interactions. Indeed, since all viruses depend on a host for virion production, a link between the isotopic compositions of hosts and the virions they produce is expected. However, stable isotope probing applied to viral DNA has never been evaluated on simple biological models. Here, this method was tested on the bacteriophage T4 and its host *Escherichia coli*. To validate that E. coli cells cultivated using a substrate enriched in ^13^C isotope were resulting on the production of 13C-labeled T4 DNA.

T4 DNA buoyant density in CsCl gradient was overall higher than the values predicted by a previously established empirical model, highlighting the need to adapt this type of models when analysing modified viral DNA. Moreover, our results show a strong correlation between the proportion of ^13^C_6_-D-glucose in the substrate used for host growth and the buoyant density of T4 DNA, validating the use of DNA SIP in viral ecology, to identify viruses infecting hosts with a specific metabolism.

## Introduction

Stable Isotope Probing (SIP) (1) has been recently employed in viral ecology to identify viruses infecting hosts with specific metabolisms (*e*.*g*. (2–7)), such as ammonia oxidation (6), methane oxidation (3) or methanogenesis (7), and it is emerging as a promising approach. Indeed, since all viruses depend on a host for virion production, a link between the isotopic compositions of hosts and the virions they produce is expected. This has been previously shown by NanoSIMS on simple biological models (5). By contrast, DNA SIP for the study of host-virus interactions has been directly applied to microbial communities (2-4, 6), without being validated on simple biological models. Moreover, either the total DNA (3, 6) or the cellular DNA (7) has been used, likely due to the difficulty of obtaining sufficient amounts of viral DNA. In principle, the application of SIP to viral DNA should be possible if enough material is obtained. Here, we validate the viral DNA SIP using T4 bacteriophage DNA obtained by propagation on *Escherichia coli* cells.

## Materials and methods

### Culture conditions and virion preparation

*Escherichia coli* strain B (DSM 613) cells were grown during 30 h at 37°C, under 400 rpm agitation, in 20 mL of M9 liquid minimal medium [M9, Minimal salts, 5X (Sigma-Aldrich), MgSO_4_ (1 mM), CaCl_2_ (0.3 mM) and D-glucose (10 % m:v)], inoculated with 20 μL of an overnight culture in LB medium (Fisher Bioragents, 25 g/L). The D-glucose was a mix of ^13^C labelled (D-Glucose-13C6, 99 % 13C, Cortecnet) and non-labelled (Sigma-Aldrich) D-glucose in various proportions. For infection with T4 bacteriophage (DSM 4505), the same conditions were employed except that the minimal medium was supplemented with 20 μL of CaCl_2_ (0.5 M) and MgCl_2_ (1 M) solutions and T4 virions were added before incubation at a multiplicity of infection of ∼10^−7^. After 30 h of culture was centrifuged at 5 000 g for 15 min at 10°C. The supernatant was filtered at a 0.22 μm pore-size with PES filters (FisherBrand). The obtained viral particle suspensions were stored at 4°C until subsequent use.

### Isotope-ratio mass spectrometry

Cells were washed once in PBS and dried overnight at 55°C. They were subsequently analysed by EA-IRMS with FlashEA 1112 Series and a Delta V Plus (ThermoFisher Scientific), as previously described (8), using 547±171 μg of cells.

### DNA extraction and dosage

Cellular DNA was extracted with the DNeasy UltraClean Microbial Kit (Qiagen), according to the instruction manual.

For extraction of T4 DNA, virions were concentrated using Amicon Ultra-15 centrifugal Filter units (Meck Millipore). For two samples only, they were concentrated by centrifugation at 20,000 g for 4 h at 4°C, followed by pellet suspension in 1 mL supernatant. The concentrated T4 suspensions were incubated with 10 μL DNase (DNase I THERMO, 1 Unite/μL) at room temperature for 20 min. The DNase was inactivated at 75°C for 5 min. Viral DNA was subsequently extracted with Phage DNA Isolation Kit (Norgen), with the following minor modifications. For proteinase K treatment (proteinase K from *Tritirachium album*, Sigma Aldrich, 20 mg/mL), samples were incubated for 15 min at 55°C with 80 μL of virion suspension. For final elution, either 2 elutions with 75 μL elution buffer, or 3 elutions with 50 μL elution buffer, were performed.

All DNAs were quantified with a Qubit fluorometer and the dsDNA HS kit (Thermo Fisher Scientific), according to the instruction manual.

### Separation of DNA in an isopycnic gradient

DNAs were separated according to their density by ultracentrifugation in an isopycnic CsCl gradient, as previously described (9).

## Results

### The ^13^C-enrichment of uninfected *E. coli* cells is consistent with the isotopic composition of the growth substrate

As a control, we first measured the isotopic composition of uninfected *E. coli* cells grown in minimal medium, by EA-IRMS (Figure 1). The sole carbon source was D-glucose containing various proportions of ^13^C_6_-D-glucose. The measured ^13^C content of cells was typically slightly inferior to the theoretical ^13^C content of the substrate, with the relative difference varying between −13.55 % and 1.40 %. Such difference cannot be explained by the inoculum, which contained LB medium, because a 1000× dilution was applied for inoculation. It could rather result from the overestimation of the ^13^C content in the ^13^C_6_-D-glucose (≥ 99 % according to the supplier), combined with biases introduced during medium preparation and IRMS measurement. Despite these minor differences, a very strong correlation was obtained between the ^13^C content of the substrate and the cells, as expected (Figure 1, linear regression, R^2^ = 0.999). Moreover, the replicates (N=2) showed high reproducibility.

**Figure 1.**
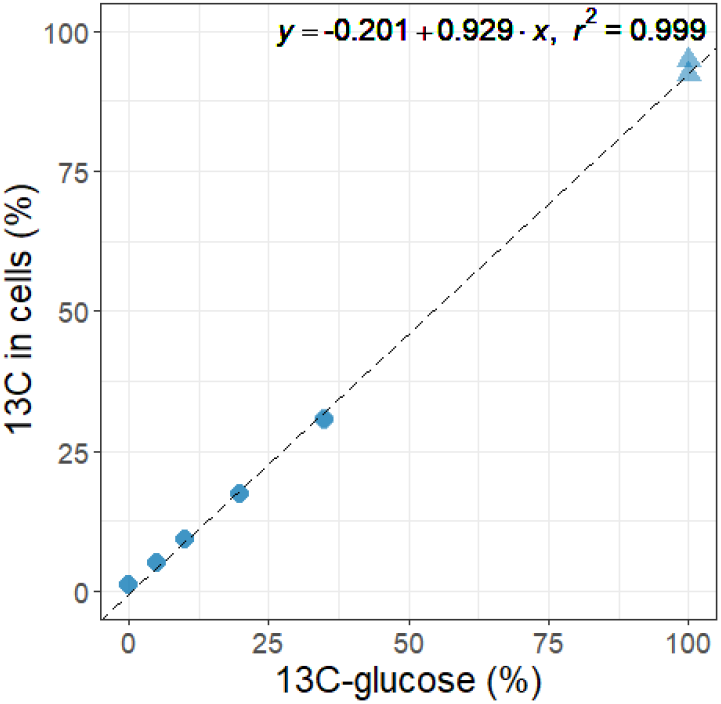
Isotopic composition of *E. coli* cells, as determined by EA-IRMS, in function of the percentage of ^13^C_6_-D-glucose employed in the D-glucose substrate (2 replicates per condition).

### The buoyant density of T4 DNA correlates with the isotopic composition of the substrate used for host growth

Subsequently, *E. coli* cells grown on minimal medium with various proportions of ^13^C_6_-D-glucose were infected by T4 bacteriophage. T4 DNA was extracted and separated on a CsCl gradient. A good reproducibility was observed among replicates, and a strong correlation was obtained between observed T4 DNA density and percentage of ^13^C_6_-D-glucose in the substrate (Figure 2A), resulting from the link between the isotopic composition of the host and the produced virions.

**Figure 2.**
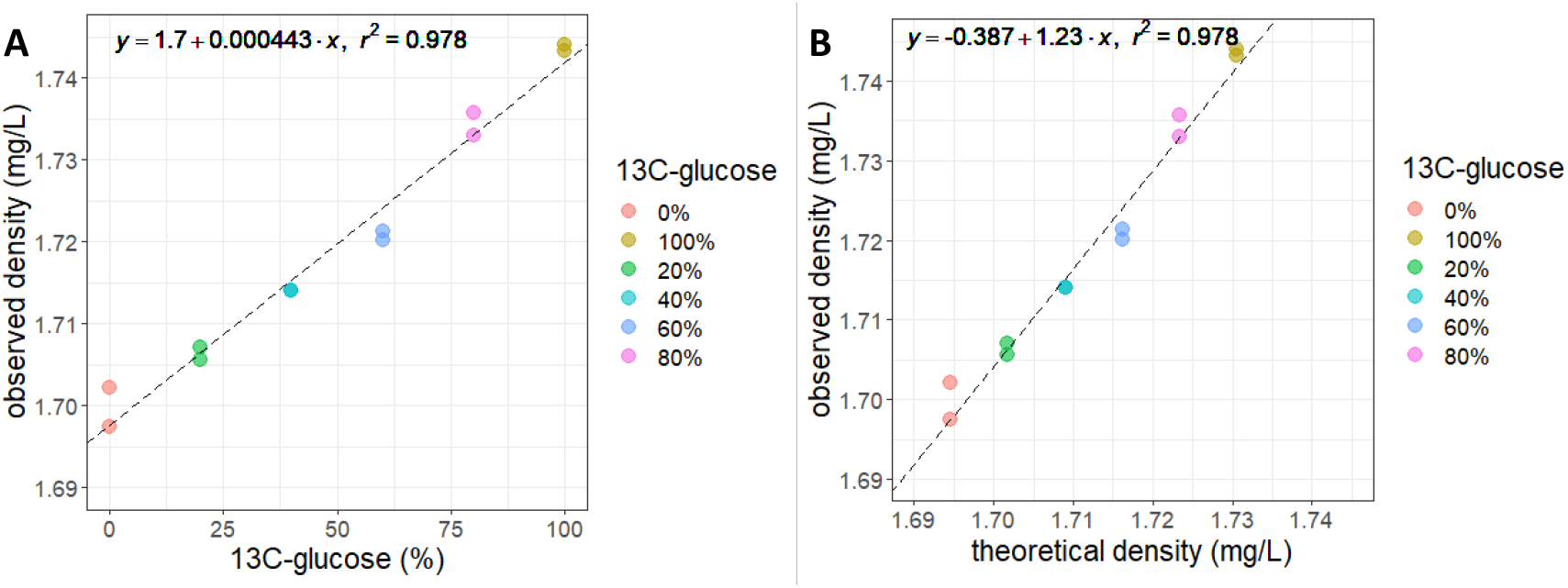
Buoyant density of T4 bacteriophage DNA measured in CsCl gradient. A. T4 DNA density as a function of the percentage of ^13^C_6_-D-glucose employed in the D-glucose substrate for host growth (2 replicates per condition). B. T4 DNA density as a function of the theoretical density calculated according to the equation presented in the text (10).

### The density of T4 DNA in CsCl gradient is overall higher than predicted according to a previously established empirical model

We calculated the theoretical expected density for T4 DNA, by relying on the previously established empirical formula (10):

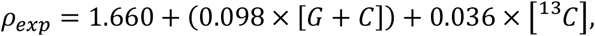

where [*G* + *C*] is the GC content of the considered DNA (0.3530 for T4, NC_000866.4), and [^13^*C*] is its ^13^C content.

The obtained predicted densities were significantly lower than the measured ones (Figure 2B). T4 DNA contains glucosylated hydroxymethylcytosine (HmC) instead of cytosine (11), affecting its buoyant density (12), so that the above formula is likely not optimal for this type of modified DNA.

## Conclusions

Our results show a strong correlation between the proportion of ^13^C_6_-D-glucose in the substrate and T4 DNA buoyant density, validating the application of DNA SIP in viral ecology. Since viral DNAs are known to be frequently modified as a mechanism to counter host defences (11), it could be worth studying their buoyant density according to their ^13^C-labeling rate, to exploit the full potential of DNA SIP profiles.

## Acknowledgements

The authors express their sincere gratitude to Olivier Tenaillon (IAME, UMR 1137, INSERM, Université Paris Cité, Paris, France) as well as Elodie Perrin and Maxime Allioux, for their assistance during the preliminary stages of this study.

This work was supported by Agence Nationale de la Recherche, France (ANR-17-CE05-0011, project VIRAME).

## Data Availability Statement

The datasets generated and analysed during the current study are available from the corresponding author on reasonable request.

